# Maternal and Neonatal Colonization with Multidrug Resistant and Extended Spectrum ß-Lactamase Producing *Escherichia coli* and *Klebsiella pneumoniae* in a Cameroonian Labour Ward

**DOI:** 10.1101/2024.02.14.579597

**Authors:** Axelle Njeuna, Luria Leslie Founou, Raspail Carrel Founou, Patrice Landry Koudoum, Aurelia Mbossi, Ariel Blocker, Stephen D. Bentley, Lucien Etame Sone

**Author notes:** **Corresponding author:** Dr Luria Leslie Founou ReMARCH CEDBCAM-RI, Yaoundé, Cameroon.

## Abstract

**Background:** *Escherichia coli* and *Klebsiella pneumoniae* rank among the primary bacterial culprits in neonatal infections and fatalities in sub-Saharan Africa. This study sought to characterize the phenotypic and genotypic features of *Escherichia coli* and *Klebsiella pneumoniae* in a labour ward in Yaoundé, Cameroon.

**Methods:** A prospective and cross-sectional study spanning five months, from February 21 to June 30, 2022. Recto-vaginal swabs were obtained from expectant mothers, and nasopharyngeal swabs were collected from their babies. The samples were cultured on eosin methylene blue agar and isolates identified using the Enterosystem 18R kit. Extended-spectrum ß-lactamase (ESBL) production was assessed using CHROMAgar ESBL™ and the double disc synergy test. Antibiotic susceptibility was determined by the Kirby-Bauer disk diffusion method. Polymerase chain reaction (PCR) was employed to detect ß-lactamase genes *bla_SHV_*, *bla_CTX_*_-*M*_ and *bla_TEM_*. ERIC-PCR was used to assess the clonal relatedness of isolates.

**Results:** *E. coli* was predominantly found in pregnant women (81%) and neonates (55%) while *K. pneumoniae* predominated in healthcare workers. Almost all pregnant women (90%) were colonized by one or more multi-drug resistant (MDR) isolates with 52% being concomitantly ESBL producers. Altogether, 22 neonates were positive for *E. coli* and/or *K. pneumoniae* and 19 (91%) were colonized by a MDR isolate. The *bla*_CTX-M_ (75%) was the leading ß-lactamase gene detected.

**Conclusion:** Our study suggests that MDR- and ESBL-*E. coli* and *K. pneumoniae* are circulating at high prevalence in labour Yaoundé. It emphasizes the necessity for strict infection prevention and control measures in conjunction with effective antimicrobial stewardship in the country.

## INTRODUCTION

Despite remarkable advances made the past decades to reduce the number of neonatal deaths globally, many newborns continue to die every year mostly from preventable causes. In 2021, globally 2.3 million children died before turning one month old, of these deaths 1 million (27.5 deaths per 1000 live births) were in neonates from sub-Saharan Africa (1). In Cameroon, the neonatal mortality rate was 23.19/1000 live births in 2021 (2), greatly exceeding United Nations Sustainable Development Goal (SDG) 3.2.2 that aims for countries to have ≤12 neonatal deaths/1000 live births by 2030 (3, 4). By 2035, it is estimated that there would be an extra 49 million newborn deaths, 52 million stillbirths, 5 million maternal deaths, and 99 million children with disabilities if significant efforts are not made (8). Therefore, continuing to reduce newborn mortality is crucial to improving infant survival. Prioritizing research appropriately and creating plans for infection prevention and control measures require a deeper understanding of the aetiology and transmission mechanisms of neonatal infections.

Neonatal sepsis remains an important cause of neonatal mortality worldwide, particularly in sub-Saharan Africa (5). The escalation of antibiotic resistance further exacerbates this problem (6, 7). Multi-drug resistant (MDR) bacteria, especially extended-spectrum β-lactamase producing *Enterobacterales* (ESBL-E) are pathogens of critical priority for research and among the leading causes of neonatal infections especially in pre-term infants (6, 7). A systematic review revealed that *Escherichia coli* and *Klebsiella pneumoniae* were among the most frequent causes of bacterial neonatal sepsis in sub-Saharan Africa with 66% of neonatal sepsis and meningitis cases caused by antibiotic-resistant *E. coli* and *K. pneumoniae* between 2008 and 2018 in the region (5). Neonates can be exposed to ESBL-E before delivery (after rupture of the amniotic membrane) or during delivery, establishing maternal rectovaginal ESBL-E carriage as an important predictor of neonatal carriage/infection (8, 9). Consequently, neonatal bacterial infections, primarily acquired at the time of delivery through maternal-neonatal transmission, are a leading preventable cause of morbidity and mortality (10).

Notwithstanding, there are currently no policies implemented for routine prenatal screening of antibiotic-resistant bacteria in Cameroon. Moreover, antibiotic usage is not well-regulated, and there is a dearth of information regarding the burden of MDR- and ESBL-*E. coli* and *K. pneumoniae* circulating strains in mother-neonate dyads in the country. Finally, the role of maternal carriage of resistant bacteria remains unknown in Cameroon, and it is unclear to what extent it could contribute to neonatal infection in the country. To the best of our knowledge, no study focusing on the burden, phenotypic and genetic profiles as well as transmission of antibiotic resistant *E. coli* and *K. pneumoniae* in mother-neonate dyads has been carried out so far in Cameroon. This study therefore ascertains the phenotypic and genotypic diversity of *E. coli* and *K. pneumoniae* in a labour ward of a hospital in Yaoundé with a view to generating knowledge and informing evidence-based strategies for better management of maternal and neonatal infections.

## MATERIALS AND METHODS

This study is reported following the Strengthening the Reporting of Observational Studies in Epidemiology for Newborn Infection (STROBE-NI) (11).

### 1. Study design and study site

A cross-sectional, prospective, and analytical study was conducted over a period of five months from February to June 2022. Sample collection was carried out during five weeks (from February 21 to March 25, 2022) in a confessional hospital in Yaoundé specializing in the care of pregnant women from pregnancy to delivery. This hospital provides antenatal care services for 700 pregnant women and practices around 450 deliveries per month.

### 2. Study population

The primary study population consisted of pregnant women with a gestational age above 32 weeks who attended the labour room of the selected hospital for delivery regardless of age, ethnicity, or HIV status. Mothers who were mentally unstable or engaged in health prognostic were excluded from the study. The secondary population was the babies of the included pregnant women. However, babies with a poor health prognosis were excluded from the study. Healthcare workers working or visiting the maternity ward were also collected. The hospital environment of the maternity ward, delivery room, and postpartum room was also considered in this study.

### 3. Recruitment and sample collection

Due to technical constraints, a saturation of sampling was implemented during the five-week sample collection period. All mothers and HCWs who met the inclusion criteria provided oral and written informed consent. A questionnaire was administered, and socio-demographic and clinical data were collected using Epicollect® 5 software (Centre for Genomic Pathogen Surveillance, Oxford, UK).

Samples were collected using sterile Amies swabs. Recto-vaginal swabs were collected from pregnant women prior to delivery, while nasopharyngeal swabs were taken from babies right after birth or before their first bath (less than 24 hours). The hands of the healthcare workers were also swabbed, as were predefined hospital environment sites (Table S1). All samples were transported within 12 hours to the Research Institute of the Centre of Expertise and Biological Diagnostics of Cameroon (<u>CEDBCAM-RI</u>).

### 4. Ethical considerations

A research permit was granted from the Ministry of Scientific Research and Innovation (N°0022/MINRESI/B00/C00/C10/C13) prior to the implementation of the study. This research was approved by the National Ethics Committee for Human Health Research (No. 2021/07/1386/CE/CNERSH/SP) and the Ethical Committee of the University of Douala (No. 3190 CEI-UDo/06/2022/M). Oral and written informed consent to participate in this study was provided by participants or the legal guardian or nearest relative of the babies. Participants were anonymized and theirand their information encoded before analysis to ensure confidentiality.

## 5. Laboratory analysis

### 5.1. Bacterial isolation and identification

All samples were cultured on Eosin-Methylene Blue (EMB) agar and incubated in a bacteriological incubator at 37°C for 18 to 24 hours. After incubation, all growing colonies were phenotypically identified with the Enterosystem 18R kit as per the manufacturer’s instructions. The screening of ESBL-producing isolates was performed with the chromogenic medium CHROMAgar ESBL™ (CHROMAgar, Paris, France).

### 5.2. Antimicrobial Susceptibility Testing

The Kirby-Bauer disk diffusion method was used to evaluate the susceptibility of E. coli and K. pneumoniae isolates against a panel of 12 antibiotics (Oxoid®), including amoxicillin-clavulanic acid (AMC, 30 µg), cefuroxime (CXM, 30 µg), ceftazidime (CAZ, 30 µg), cefotaxime (CTX, 30 µg), cefepime (FEP, 30 µg), cefoxitin (FOX, 30 µg), amikacin (AK, 30 µg), gentamicin (GEN, 30 µg), ciprofloxacin (CIP, 5 µg), chloramphenicol (C, 30 µg), doxycycline (DOX, 30 µg), and trimethoprim-sulfamethoxazole (TMP/SXT, 25µg). For all antibiotics, the EUCAST (12) clinical breakpoints were used for interpretation. But for doxycyclin, and trimethoprime-sulfamethoxazole, the CLSI 2020 (13) guideline was used. Multidrug resistance (MDR), which is the resistance of bacteria to at least one antibiotic from three or more families of antibiotics, was also evaluated.

### 5.3. Genotypic analysis

#### 5.3.1. Genomic amplification of *bla_SHV,_ bla_CTX-M_* and *bla_TEM_*

The genomic DNA of *E. coli* and *K. pneumoniae* isolates was extracted using a modified boiling method (14). After incubation, the suspension was centrifuged at 9500 rpm for 5 min, and then 300 µL of the supernatant having DNA was transferred to a new Eppendorf tube and stored at −30 °C for future use. The detection of the *bla_SHV_* gene was performed by conventional PCR in a 10 µL reaction mixture consisting of 5 µL of DreamTaq™ Green Polymerase 2X (ThermoFisher Scientific™, Vilnius, Lithuana), 2.8 µL of nuclease-free water, 0.1 µL of each forward and reverse primer [10 µM], and 2 µL of template DNA. Likewise, detection of *bla_CTX-M_* and *bla_TEM_*genes was performed by multiplex-PCR (M-PCR) in a 10 µL reaction mixture consisting of 5 µL of DreamTaq Green Polymerase 2X (ThermoFisher Scientific™, Vilnius, Lithuana), with 0.1 µL of each forwad (CTX-Mu1 and TEM-F) and reverse (CTX-Mu2 and TEM-R) primers [50 µM] and 2 µL of DNA. Primer sequences previously reported [9], were all synthesized by Inqaba Biotec West Africa. All amplification reactions took place in a BIO-RAD T100 thermal cycler (Bio-Rad Laboratories, Marnes-la-Coquette, France) following the programming conditions described previously (14).

#### 5.3.2. Agarose gel electrophoresis and DNA visualization

After amplification, DNA electrophoresis was performed on a 1.5% (wt/vol) agarose gel run at 90 V for 45 min with a molecular ladder of 100 bp (New England Biolabs, MA, USA). The gel was then stained in an ethidium bromide solution (0.5 µg/mL) for 15 min and briefly unstained with water. The amplicons were visualized under UV light using a G-BOX Chemi XL gel documentation system (Syngene, Cambridge, UK). Figure 1 shows the visualization of amplicons from various samples with the targeted ESBL genes.

**Figure 1:**
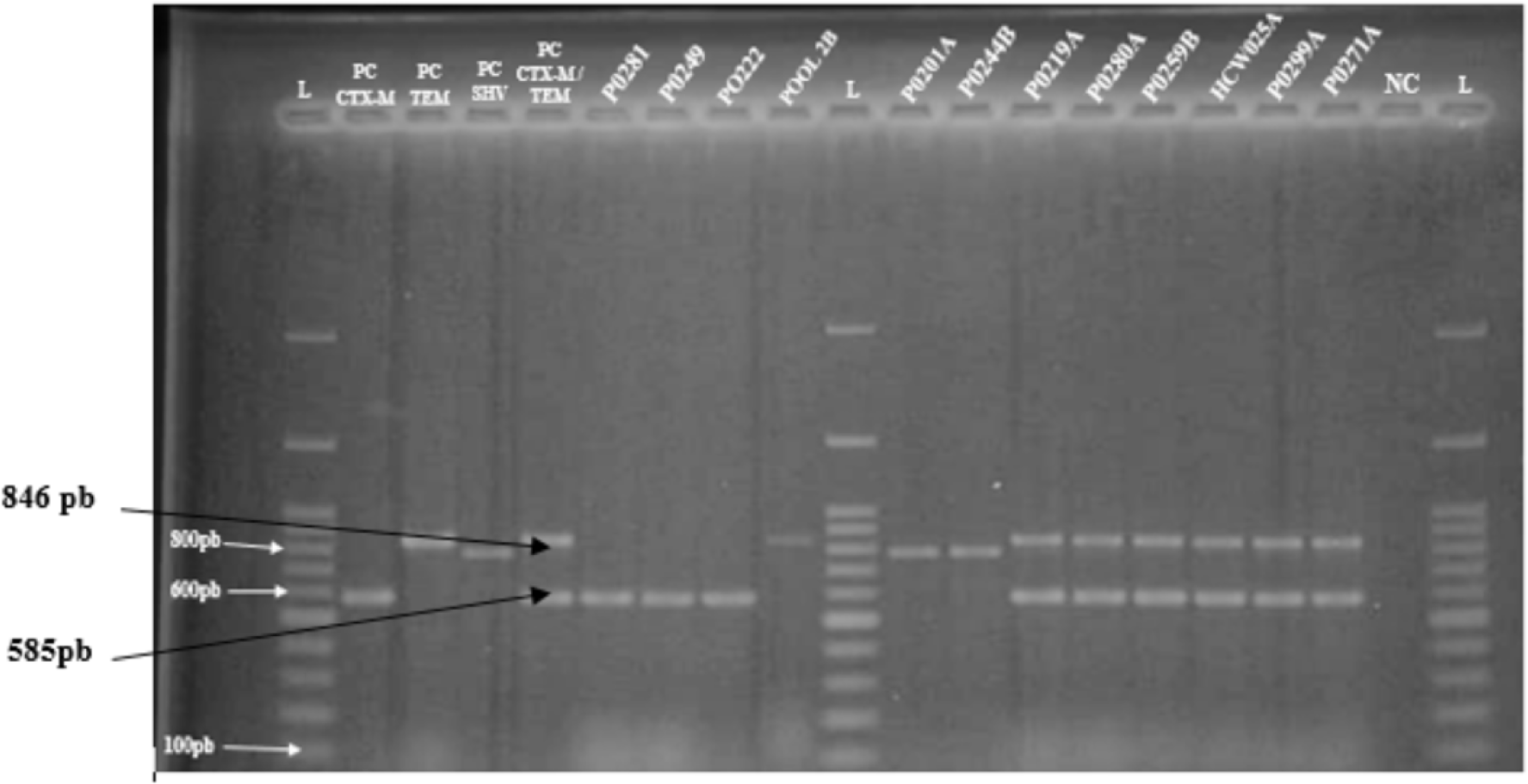
Agarose gel (1.5%) electrophoresis of PCR amplified genes of selected isolates. L: 100 bp Ladder; PC: Positive control; NC: Negative control; Isolates: P0281, P0249, P0222, POOL 2B, P0201A, P0244B, P0219A, P0280A, P0259A, HCW025A, P0229A, PO271A.

#### 5.3.3. Genotypic relatedness

Enterobacterial Repetitive Intergenic Consensus-Polymerase Chain Reaction (ERIC-PCR) was used to show the clonal relatedness between the isolates originating from mother-baby pairs, HCWs, and the environment. The primers ERIC1 5’ATGTAAGCTCCTGGGGATTCAC3’ and ERIC2 5’AAGTAAGTGAC TGGGGTGAGCG 3’ [10] were used, and ERIC-PCR reactions were performed in a 10 μl volume containing 5 μl of DreamTaq Green Polymerase 2X (ThermoFisher Scientific™, Vilnius, Lithuana), 0.1 μl of each primer, 2.8 μl of nuclease-free millipure water, and 2 μl of DNA template. The amplification steps were as previously described [10]. The generated PCR products were resolved by horizontal electrophoresis on 1.5% (wt/vol) Tris-Acetate-EDTA agarose gels (Merck, Germany) with Quick-load®1-kb (Biolabs, New England) and run in a 110 V electric field for 2 h 30 min. Electrophoresis gels were visualized by a UV light transilluminator. Gel images were exported to GelJ software (version 2.0) for cluster analyses. Metadata, phenotypic, and genotypic data were incorporated into the dendrograms for better clonal relatedness inference.

### 6. Quality controls

*E. coli* ATCC 35218 and *Klebsiella pneumoniae* ATCC 700603 strains were used as positive controls for the *bla_TEM_* and *bla_SHV_* genes, respectively. For the *bla_CTX-M_* gene, a previously whole genome sequenced *E. coli* isolate (unpublished result) served as an internal quality control. *E. coli* ATCC 35218 and *Klebsiella pneumoniae* ATCC 700603 strains were used as positive controls for the *bla_TEM_* and *bla_SHV_* genes, respectively. For the *bla_CTX-M_* gene, a previously whole genome-sequenced *E. coli* isolate (unpublished result) served as an internal quality control.

### 7. Data management and statistical analysis

Data collected in EpiCollect® were exported to Microsoft Office Excel 2016 (Microsoft® Excel). Data cleaning and data analysis were performed using R software (version 4.1.0) and RStudio (version 2021.09.0). A participant was considered positive to *E. coli* or *K. pneumoniae* when one colony of any of the species was detected in the samples. Likewise, a participant was ESBL positive when at least one ESBL colony was detected. A participant was considered multidrug resistant positive when an *E. coli* or *K. pneumoniae* isolate showed resistance to at least three antibiotics from three or more families of antibiotics, with or without the presence of an ESBL phenotype. The Fisher exact and Chi-square test (where appropriate) were used to compare the proportions among categorical variables. Results were considered statistically significant at a *p*-value < 0.05. Missing data were not considered in the analysis.

## RESULTS

### 1. Population socio-demographic and clinical characteristics in relation to colonization status

### 1.1. Pregnant women

Out of the 103 women contacted, 100 were enrolled and 93 provided samples. The mean age of the included participants was 27.8 years (± std. 5.6 years), and all lived in urban areas. The average gestational age of pregnant women was 38.9 weeks (±std. 4.4 weeks). Among the 93 women sampled, 87% (81/93) were positive for *E. coli* and/or *K. pneumoniae* (Table 1). Of these, 90% (73/81) were colonized by at least one MDR isolate with 52% (42/81) of these being concomitantly colonized by at least one ESBL isolate (Figure 2).

**Figure 2.**
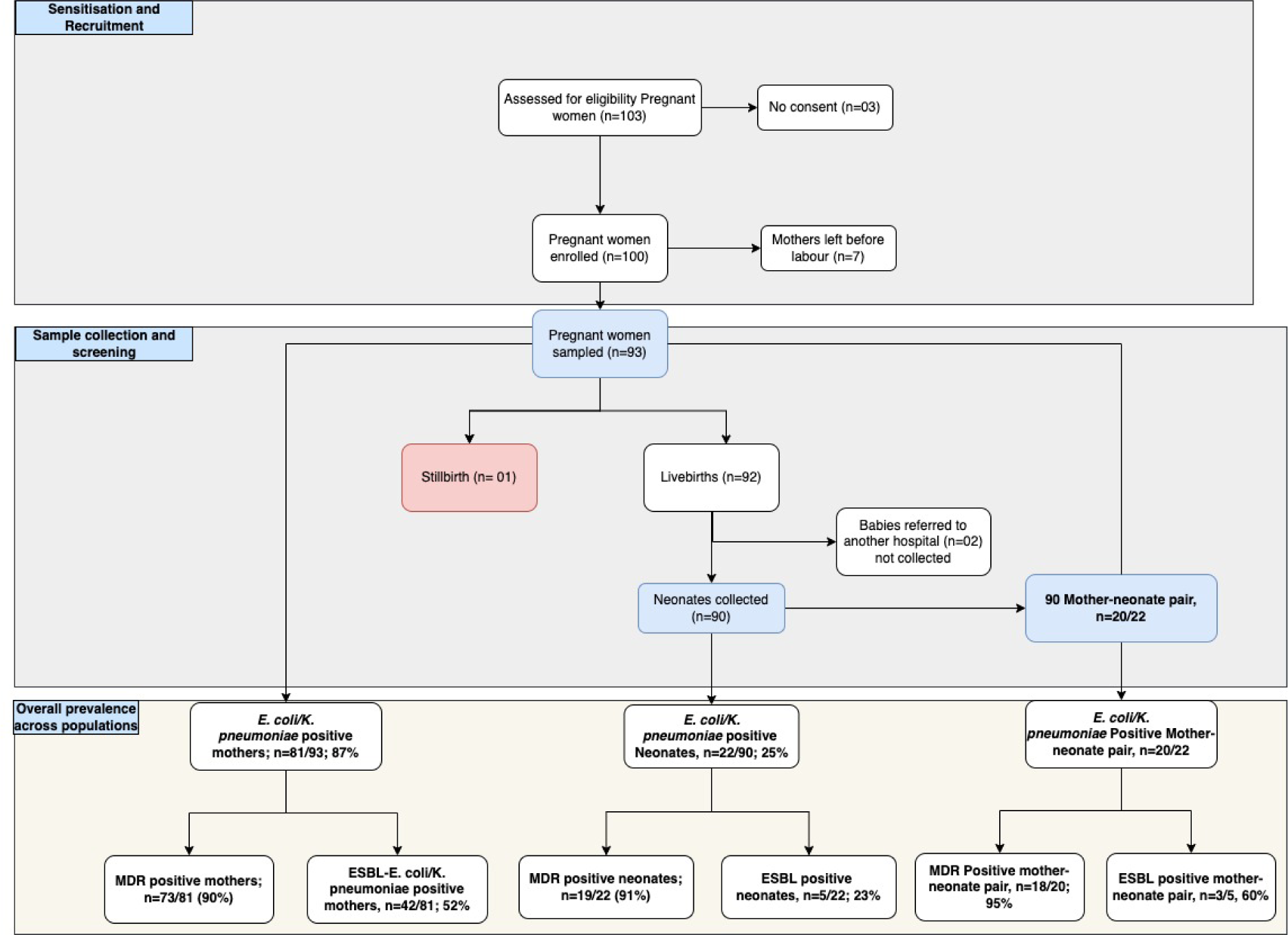
Flow chart summarizing mothers’ and neonates’ recruitment, screening and colonization status.

**Table 1.** Distribution of socio-demographic and clinical characteristics among pregnant women in relation with the colonization of *E. coli* and/or *K. pneumoniae*.

| Variables |  | Positive*,<br>n (%) | p-value | MDR<br>positive<br>n (%) | p-value | ESBL<br>positive<br>n (%) | p-value |
| --- | --- | --- | --- | --- | --- | --- | --- |
| Overall (N= 93) |  | 81(87.09) | - | 73 (90.12) | - | 42(45.16) | - |
| Age (years) | [15 -24] | 25 (30.86) | 0.9972 | 21 (31.82) | 0.155 | 15 (35.71) | 0.3984 |
|  | [25-34] | 46 (56.79) |  | 35 (53.03) |  | 19 (45.25) |  |
|  | [35-44] | 10 (12.35) |  | 10 (15.15) |  | 8 (1.04) |  |
| Residence | Urban | 81 (100) | / | 73 (100) | / | 42 (100.00) | / |
| Education | Primary | 3 (3.70) | 0.2315 | 3 (4.11) | 0.785 | 2 (4.76) | 0.4129 |
|  | Secondary | 53 (65.43) |  | 48 (65.75) |  | 30 (71.43) |  |
|  | High school | 25 (30.86) |  | 22 (30.14) |  | 10 (23.81) |  |
| Person living at home | [1-7] | 67 (8.71) | 0.4272 | 52 (78.79) | 0.045 | 34 (80.96) | 0.0294 |
|  | [8-14] | 13 (16.04) |  | 13 (19.69) |  | 8 (19.04) |  |
|  | [15-21] | 1 (1.25) |  | 1 (1.52) |  | 0 |  |
| Occupation | Others | 23 (28.40) | 0.7252 | 20 (27.40) | 1.000 | 11 (26.19) | 0.3253 |
|  | Seller | 3 (3.70) |  | 3 (4.11) |  | 3 (7.14) |  |
|  | Student | 26 (32.10) |  | 23 (31.51) |  | 15 (35.71) |  |
|  | Farmer | 1 (1.23) |  | 1 (1.37) |  | 0 (0) |  |
|  | Official | 6 (7.41) |  | 6 (8.22) |  | 4 (9.52) |  |
|  | housewife | 20 (24.69) |  | 18 (24.66) |  | 9 (21.43) |  |
|  | Healthcare workers | 2 (2.47) |  | 2 (2.74) |  | 0 (0) |  |
| Average monthly income (CFA Francs) | < 30.000 | 45 (55.56) | 0.8633 | 40 (54.79) | 1.000 | 25 (59.52) | 0.2257 |
|  | 30.000– 60.000 | 16 (19.75) |  | 14 (19.18) |  | 10 (23.81) |  |
|  | 60.000– 90.000 | 14 (17.28) |  | 13 (17.81) |  | 5 (11.90) |  |
|  | 90.000– 120.000 | 2 (2.47) |  | 2 (2.74) |  | 1 (2.38) |  |
|  | 120.000– 150.000 | 3 (3.70) |  | 3 (4.11) |  | 0 (0) |  |
|  | > 150. 000 | 1 (1.23) |  | 1 (1.37) |  | 1 (2.38) |  |
| Animal contact | No | 61 (75.31) | 0.2860 | 54 (73.97) | 0.672 | 31 (73.81) | 0.7454 |
|  | Yes | 20 (24.69) |  | 19 (26.03) |  | 11 (26.19) |  |
| Drinking water | Borehold | 52 (64.20) | 0.0074 | 48 (65.75) | 0.100 | 29 (69.05) | 0.5346 |
|  | Mineral | 5 (6.17) |  | 3 (4.11) |  | 1 (2.38) |  |
|  | Tap water | 18 (22.22) |  | 17 (23.29) |  | 9 (21.43) |  |
|  | Source | 6 (7.41) |  | 5 (6.85) |  | 3 (7.14) |  |

**Table 1. Continued**
| Variable | | Positive*<br>n (%) | P-value | MDR positive*&<br>n (%) | P-value | ESBL positive\$<br>n (%) | P-value |
| --- | --- | --- | --- | --- | --- | --- | --- |
| Hospitalisation | No | 75 (92.59) | 0.28 | 69 (94.52) | 0.20 | 37 (88.10) | 0.20 |
|  | Yes | 6 (7.41) |  | 4 (5.48) |  | 5 (11.90) |  |
| Antibiotic use | No | 76 (93.83) | 0.22 | 68 (93.15) | 0.67 | 40 (95.24) | 0.66 |
|  | Yes | 5 (6.17) |  | 5 (6.85) |  | 2 (4.76) |  |
| Status to infectious diseases | VIH | 2 (2.47) | 0.63 | 2 (2.74) | 0.54 | 1 (2.38) | 0.54 |
|  | Hepatite B | 2 (2.47) |  | 2 (2.74) |  | 2 (4.76) |  |
|  | Chlamydia | 1 (1.23) |  | 1 (1.32) |  | 0 (0) |  |
|  | None | 76(93.83) |  | 68 (93.15) |  | 39 (92.86) |  |
| Gravidity | Primipara | 33 (40.74) | 1.00 | 30 (41.10) | 0.62 | 16 (38.10) | 0.62 |
|  | Multipara | 48 (59.26) |  | 43 (58.90) |  | 26 (61.90) |  |
| Death in utero | No | 80 (98.77) | 1.00 | 72 (98.63) | 0.48 | 42 (100.00) | 0.48 |
|  | Yes | 1 (1.23) |  | 1 (1.37) |  | 0 (0) |  |
| Abortion | No | 51 (62.96) | 0.53 | 44 (60.27) | 0.11 | 23 (54.76) | 0.11 |
|  | Yes | 30 (37.04) |  | 29 (39.73) |  | 19 (45.24) |  |
| Stillbirth | No | 73 (90.12) | 1.00 | 68 (93.15) | 0.03 | 37 (88.10) | 0.71 |
|  | Yes | 8 (9.88) |  | 5 (6.85) |  | 5 (11.90) |  |
| Gestational age | Min / Max | 0 / 42.0 | 0.77 | 0 / 42.0 | 0.93 | 32.0 / 42.0 | 0.93 |
|  | Med [IQR] | 40.0 [38.0;40.0] |  | 39.0 [38.0;40.0] |  | 40.0 [38.0;40.0] |  |
|  | Mean (std) | 38.8 (4.7) |  | 38.7 (4.9) |  | 39.2 (2.0) |  |
| Antenatal care visit | Min / Max | 0/9.0 | 0.54 | 0/9.0 | 0.66 | 0/8.0 | 0.25 |
|  | Med [IQR] | 4.0 [3.0;5.0] |  | 4.0 [3.0; 5.0] |  | 4.0 [3.0;5.0] |  |
|  | Mean (std) | 4.1 (1.7) |  | 4.1 (1.7) |  | 3.8 (1.6) |  |
| No. birth | Min / Max | 1.0/5.0 | 0.31 | 1.0/5.0 | 0.83 | 1.0/5.0 | 0.01 |
|  | Med [IQR] | 2.5[1.0;3.2] |  | 3.0[1.5;3.5] |  | 3.0[2.0;4.0] |  |
|  | Mean (std) | 2.6 (1.3) |  | 2.6 (1.3) |  | 3.0 (1.3) |  |

\*Colonization by *E. coli* and/or *K. pneumoniae*, &Colonization by MDR-*E. coli* and/or MDR-*K. pneumoniae*, \$Colonization by ESBL-*E. coli* and/or ESBL-*K. pneumoniae*, ESBL: Extended spectrum β-lactamase, MDR: Multidrug resistance

### 1.2. Neonates

From the 93 mothers enrolled, 90 neonates were included and sampled. Most of them were male (57%, 53/93) and the average weight was 3290 g (±std 644.88 g). Out of these, 25% (22/90) were positive for *E. coli* and/or *K. pneumoniae* with male being more positive than female (64% vs 36%, p=0.47) although without statistical significance. Most of the positive neonates weighted ≥2500 g (96%, 21/22) and had a median APGAR score of 9.0 (IQR:8.0-10.0) at birth. In addition, 91% (19/22) neonates were colonized by MDR positive and 23% (5/22) were ESBL positive (Table 2, Figure 2).

**Table 2.** Colonization status in relation to socio-demographic and clinical characteristics of neonates.

| Variable |  | Positive*<br>n (%) | P- value | MDR positive<br>n (%) | P- value | ESBL positive<br>n (%) | P- value |
| --- | --- | --- | --- | --- | --- | --- | --- |
| Overall (N= 90) |  | 22<br>(24.44) | - | 20(90.91) | - | 5 (22.73) | - |
| Sex | Male | 14 (63.64) | 0.47 | 12 (60) | 0.52* | 2 (40.00) | 0.31 |
|  | Female | 8 (36.36) |  | 8 (40) |  | 3 (60.00) |  |
| Gestational age | Min / Max | 32.0 / 42.0 | 0.64 | 32.0 / 42.0 | 0.41 | 32.0 / 40.0 | 0.20 |
|  | Med [IQR] | 39.5<br>[38.0;40.0] |  | 39.0 [38.0;40.0] |  | 39.0 [36.0;39.0] |  |
|  | Mean (std) | 39.0 (2.2) |  | 38.9 (2.3) |  | 37.2 (3.3) |  |
| Weight | < 2500g | 1(4.54) | 0.90 | 1(7.14) | 0.33 | 0(0) | 0.5907 |
|  | ≥2500g | 21(95.46) |  | 13(92.86) |  | 5(100) |  |
| APGAR score at birth | Min / Max | 0 / 10.0 | 0.1 | 0 / 10.0 | 0.40 | 9.0 / 10.0 | 0.0396 |
|  | Med [IQR] | 9.0 [8.0;10.0] |  | 9.0 [8.0;10.0] |  | 10.0 [10.0;10.0] |  |
|  | Mean (std) | 8.6 (2.2) |  | 8.7 (2.3) |  | 9.8 (0.4) |  |
| APGAR score after 5 min | Min / Max | 0 / 10.0 | 0.98 | 0 / 10.0 | 0.57 | 10.0 / 10.0 | 0.3245 |
|  | Med [IQR] | 10.0<br>[10.0;10.0] |  | 10.0 [10.0;10.0] |  | 10.0 [10.0;10.0] |  |
|  | Mean (std) | 9.5 (2.1) |  | 9.4 (2.2) |  | 10.0 (0) |  |
| *E. coli* or *K. pneumoniae*<br>Positive mother | No | 2 (9.09) | 0.72 | 1 (5) | 0.18 | 0 (0) | 1.00 |
|  | Yes | 20 (90.91) |  | 19 (95) |  | 5 (100.00) |  |
| ESBL status of mother | No | 8 (40) | 0.36# | 8 (42.11) | 1.00† | 2 (40) | 1.00 |
|  | Yes | 12(60) |  | 11 (57.89) |  | 3 (60) |  |
| MDR status of mother | No | 1 (5) | 0.67# | 1 (5.26) | 1.00† | 0 | 1.00 |
|  | Yes | 19 (95) |  | 18 (94.74) |  | 5 (100) |  |

\*Colonization by *E. coli* and/or *K. pneumoniae*, &Colonization by MDR-*E. coli* and/or MDR-*K. pneumoniae*, \$Colonization by ESBL-*E. coli* and/or ESBL-*K. pneumoniae*, ESBL: Extended spectrum β-lactamase, MDR: Multidrug resistance, #Missing values (n=02) were not included in the analysis, †Missing values (n=01) were not included in the analysis, Med: Median, IQR: Interquartile range, Std: Standard deviation, Min: minimum, Max: Maximum

### 1.3. Healthcare workers

Twenty-five workers were enrolled in the study, with a 100% response rate. Table 3 presents the different factors associated with *E. coli* and *K. pneumoniae* colonization in this population. Five healthcare workers were positive for *E. coli* and/or *K. pneumoniae*. Many workers colonized with *E. coli* and/or *K. pneumoniae* lived in the district of Yaoundé V (40%); and were attached to the maternity ward (60%). Most admitted washing their hands five to ten times a day (80%; 4/5) and only after any act on the mother or neonates (Table 3).

**Table 3.** Distribution of colonization status in relation to socio-demographic characteristics of healthcare workers.

| Variable |  | Positive*<br>n (%) | P-value | ESBL<br>positive<br>n (%) | P-value | MDR<br>positive,<br>n (%) | P- value |
| --- | --- | --- | --- | --- | --- | --- | --- |
| **Overall (N= 25)** |  | **5 (20)** | **-** | **3 (60)** | **-** | **5 (100)** | **-** |
| **Age** | Min / Max | 1.0 / 15.0 | 0.38 | 1.0 / 15.0 | 1.00 | 1.0 / 15.0 | - |
|  | Med [IQR] | 7.0 [2.0;9.0] |  | 7.0 [4.0;11.0] |  | 7.0 [2.0;9.0] |  |
|  | Mean (std) | 6.8 (5.7) |  | 7.7 (7.0) |  | 6.8 (5.7) |  |
| **Service** | Others | 0 (0) | 0.53 | 0 (0) | 0.40 | 0 (0) | - |
|  | Operating room | 2 (40) |  | 2 (66.67) |  | 2 (40) |  |
|  | Consultation | 0 (0) |  | 0 (0) |  | 0 (0) |  |
|  | Hospitalisation | 0 (0) |  | 0 (0) |  | 0 (0) |  |
|  | Maternity | 3 (60) |  | 1 (33.33) |  | 3 (60) |  |
| **Animals** | No | 4 (80) | 0.34 | 2 (66.67) | 1.00 | 4 (80) | - |
|  | Yes | 1 (20) |  | 1 (33.33) |  | 1 (20) |  |
| **Frequency of**<br>**hand washing**<br>**per day** | 1 - 5 times per day | 0 (0) | 0.36 | 0 (0) | 1.00 | 0 (0) | - |
|  | 5 - 10 times per day | 4 (80) |  | 2 (66.67) |  | 4 (80) |  |
|  | 10 - 30 times per day | 0 (0) |  | 0 (0) |  | 0 (0) |  |
|  | more than 30 times | 1 (20) |  | 1 (33.33) |  | 1 (20) |  |
| **Time to wash**<br>**your hands** | After any procedure<br>on a pregnant woman<br>or newborn | 4 (80) | 0.36 | 2 (66,66) | 1.00 | 4 (80) | - |
|  | Before and after any<br>procedure in a pregnant<br>woman or newborn | 1 (20) |  | 1 (33,33) |  | 1(20) |  |
| **Residence**<br>**District** | YAOUNDE 1 | 0 (0) | 0.11 | 0 (0) | 1.00 | 0 (0) | - |
|  | YAOUNDE 2 | 1 (20) |  | 0 (0) |  | 1 (20) |  |
|  | YAOUNDE 3 | 0 (0) |  | 0 (0) |  | 0 (0) |  |
|  | YAOUNDE 4 | 1 (20) |  | 1 (33.33) |  | 1 (20) |  |
|  | YAOUNDE 5 | 2 (40) |  | 1 (33.33) |  | 2 (40) |  |
|  | YAOUNDE 6 | 1 (20) |  | 1 (33.33) |  | 1 (20) |  |
|  | YAOUNDE 7 | 0 (0) |  | 0 (0) |  | 0 (0) |  |
| **Taking**<br>**medication** | No | 5 (100) | 0.54 | 3 (100) | 1.00 | 5 (100) | - |
|  | Yes | 0 (0) |  | 0 (0) |  | 0 (0) |  |
| **Knowledge of**<br>**bacterial**<br>**resistance** | No | 1 (20) | 1.00 | 1 (33.33) | 1.00 | 5 (100) | - |
|  | Yes | 4 (80) |  | 2(66.67) |  | 0(0) |  |

\*Refers to the colonization by *E. coli* and/or *K. pneumoniae*

### 2. Prevalence in mother-neonate dyads

Twenty (91%), 19 (95%), and 12 (60%) of the 22 positive neonates were born to positive, MDR-positive, and ESBL-positive mothers, respectively. Of the twenty newborns with MDR positivity, 18 (95%) were born to mothers with MDR positivity, while 11 (58%) were delivered to mothers with ESBL positivity. Interestingly, 60% (3/5) of ESBL positive neonates were born to ESBL positive mothers while all of them were delivered from MDR positive mothers although without statistical significance.

### 1. 3. Overall prevalence of *E. coli* and *K. pneumoniae* isolates

Altogether, 217 Gram-negative bacilli were isolated from the different populations, among which 56% were *E. coli* (95/170) and 44% *K. pneumoniae* (75/170) isolates. Most of these isolates, were detected in pregnant women (82%, 140/170), of which the majority were *E. coli* (55% vs 45%, p=0.01). Twenty-two (16%) isolates, including 16 (73%) *E. coli* and six (27%) *K. pneumoniae* were identified in neonates (p=0.01). Six (4%) isolates were detected from healthcare workers, including one *E. coli* isolate (17%) and five *K. pneumoniae* isolates (83%). Finally, only one isolate of each species was detected in the environment.

### 1. 4. Prevalence of *E. coli* and *K. pneumoniae* isolates in mother and neonate pairs

Among the 22 isolates detected in neonates, 20 (91%) were born to positive mothers including six (27%), four (18%), and eight mother-neonate pairs positive to *E. coli*, *K. pneumoniae* and a combination of *E. coli* and *K. pneumoniae*, respectively. Two couple of mother and neonates were positive to *E. coli* and *K. aerogenes* while two were discordant with positive neonates and negative mothers.

### 1. 5. Antimicrobial resistance profiles of *E. coli* and *K. pneumoniae* isolates

Altogether, 78% (132/170) of isolates were MDR with 51% being *E. coli* (68/132) and 49% *K. pneumoniae* (64/132). Intriguingly, all isolates of healthcare workers were MDR, while an important level of MDR-*E. coli* isolates was recorded among neonates (71%, 15/21) with strong statistical significance (p=0.0002) (Figure 3). In contrast, similar prevalences of MDR-*E. coli* (49%, 52/105) and MDR-*K. pneumoniae* (51%, 53/105) were detected in mothers. Interestingly, of the MDR isolates detected, 44% (58/132) were concomitantly ESBL producers, most of these were *K. pneumoniae* (60%; 35/58 vs 40%; 23/58) (Figure 3). MDR-ESBL-*K. pneumoniae* predominated in mothers (56%, 28/50) and neonates (100%, 5/5) with high statistical significance (p<0.0001).

**Figure 3:**
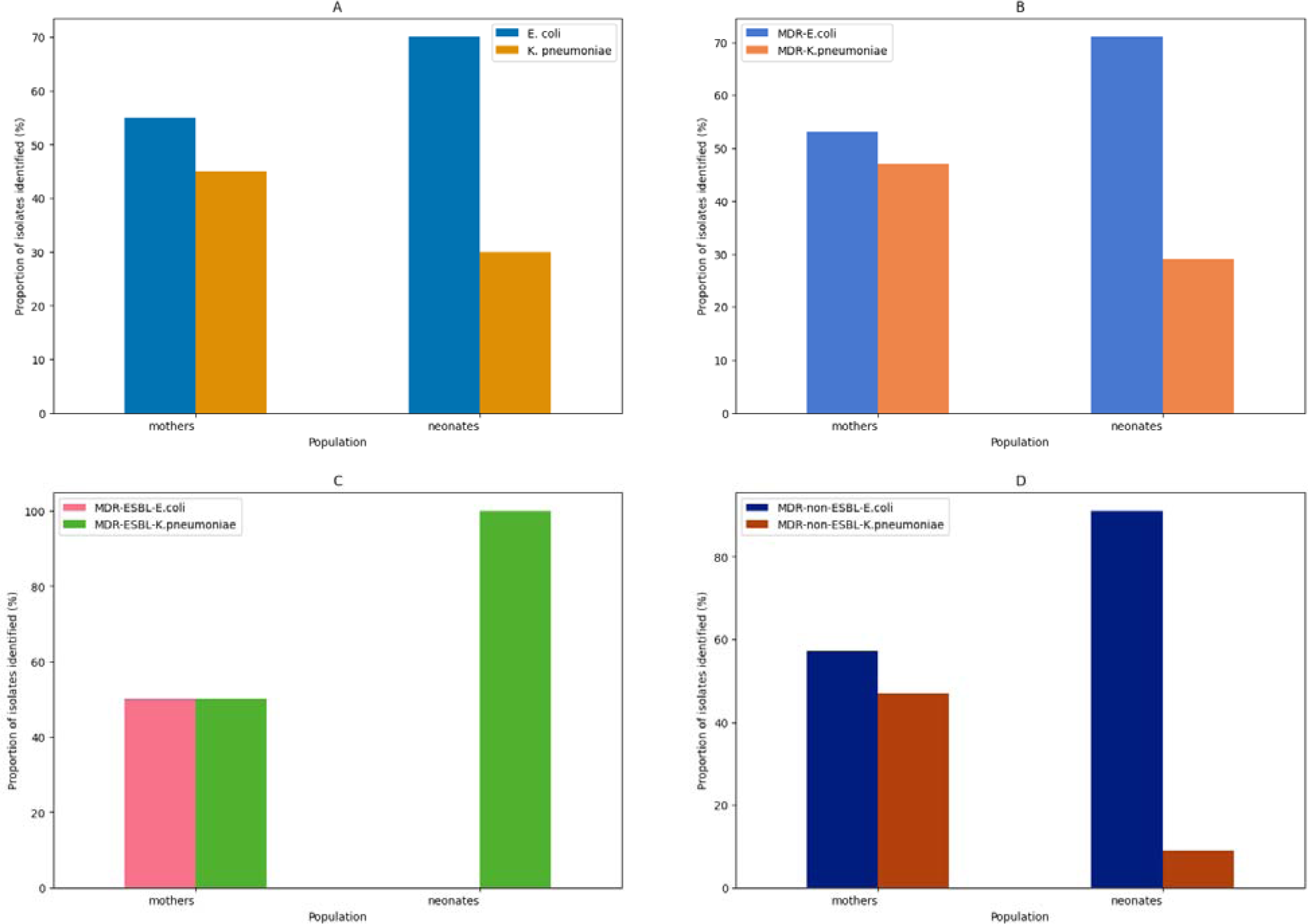
Distribution of *E. coli* and *K.* pneumoniae (A), MDR-*E. coli* and MDR-*K. pneumoniae* (b), MDR-ESBL-*E. coli* and MDR-ESBL-*K. pneumoniae* (C), and MDR-non-ESBL-*E. coli* and MDR-non-ESBL-*K. pneumoniae* (D) isolates in mothers and neonates.

Intriguingly, a total of 10/68 (15%) isolates were ESBL producers only, among which 60% (6/10) were *E. coli* and 40% (4/10) *K. pneumoniae*. Likewise, 56% (74/132) isolates were MDR-non-ESBL producers with 61% (45/74) and 39% (29/74) being MDR-non-ESBL-*E. coli* and MDR-non-ESBL-*K. pneumoniae* isolates, respectively. More specifically, 94% (15/16) of MDR-non-ESBL-*E. coli* isolates were recorded in neonates whereas 55% (30/55) of MDR-non-ESBL-*E. coli* isolates were detected in mothers with high statistical significance (p<0.0001) (Figure 3). Half (3/6, 50%) of the healthcare worker isolates were MDR-non-ESBL-*K. pneumoniae* while the sole environmental *K. pneumoniae* isolate was ESBL producer only and *E. coli* was non-MDR-non-ESBL.

Overall, isolates displayed high resistance to cotrimoxazole (83%; 141/171), doxycycline (59%; 100/171), and a low level of resistance to amikacin (6.43%; 11/171) (Table 4). *E. coli* was more resistant to cotrimoxazole (80%) and doxycycline (58.9%), while *K. pneumoniae* was more resistant to cotrimoxazole (86%).

**Table 4.** Distribution of antibiotic resistance among *E. coli* and *K. pneumoniae* isolates.

| Bacterial Species (n) | β-lactams, n (%) | | | | | | Other antibiotic families, n(%) | | | | | |
| --- | --- | --- | --- | --- | --- | --- | --- | --- | --- | --- | --- | --- |
|  | AUG | CXM | CTX | CAZ | FEP | FOX | AK | GEN | DOX | CIP | CHL | TMP/SXT |
| *E. coli* (95) | 21<br>(22.1) | 31<br>(39.7) | 30<br>(31.6) | 15<br>(15.8) | 21<br>(22.1) | 7<br>(22.1) | 6<br>(6.32) | 10<br>(10.5) | 56<br>(58.9) | 17<br>(17.9) | 21<br>(22.1) | 76<br>(80) |
| *K. pneumoniae* (75) | 43<br>(56.57) | 45<br>(59.21) | 32<br>(42.10) | 25<br>(32.89) | 28<br>(36.84) | 28<br>(36.84) | 5<br>(6.57) | 18<br>(23.68) | 44<br>(57.89) | 20<br>(26.31) | 22<br>(28.94) | 65<br>(85.52) |
| **Total (170)** | **64**<br>**(37.42)** | **76**<br>**(44.44)** | **62**<br>**(36.25)** | **40**<br>**(23.39)** | **49**<br>**(28.65)** | **35**<br>**(20.46)** | **11**<br>**(6.43)** | **28**<br>**(16.37)** | **100**<br>**(58.47)** | **37**<br>**(21.64)** | **43**<br>**(25.14)** | **141**<br>**(82.45)** |

AUG: amoxicillin – clavulanic acid; CXM: Cefuroxime; CTX : Cefotaxime CAZ : Ceftazidime ; FEP : Cefepime ; FOX : Cefoxitin; AK : Amikacin ; GEN : Gentamicin ; DOX : Doxycycline ; CIP : Ciprofloxacin ; CHL : chloramphenicol ; TMP/SXT : Trimethoprim-sulfamethoxazole (cotrimoxazole).

### 6. Prevalence of resistance genes involved in ESBL production

Of the 68 ESBL-producing isolates, the *bla_CTX-M_* (75%, 51/68) was the most frequent genes followed by *bla_TEM_* (43%, 29/68) and *bla_SHV_* (41%, 28/68) genes (Table 5). When analysed at the population level, *bla_CTX-M_* predominated in pregnant women (48/59, 81%) while all genes had a 40% prevalence among neonates. The two ESBL positive isolates from healthcare workers were harboured individually *bla_CTX-M_*and *bla_TEM_*.

**Table 5.** Prevalence of ß-lactamase resistance genes among ESBL-positive isolates according to the source of isolation.

| **SOURCE** | ***bla*<sub>CTX-M</sub>, n (%)** | ***bla*<sub>TEM</sub>, n (%)** | ***bla*<sub>SHV</sub>, n (%)** |
| --- | --- | --- | --- |
| **Overall N=68 (n)** | **51 (75)** | **29 (42.64)** | **28 (41.17)** |
| **Pregnant women (59)** | 48 (81.35) | 25 (42.37) | 26 (44.06) |
| **Newborns (5)** | 2(40) | 2(40) | 2 (40) |
| **Heathcare workers (2)** | 1 (50) | 1 (50) | 0 |
| **Environment (1)** | 0 | 1(100) | 0 |

### 7. Genomic fingerprinting by ERIC-PCR

ERIC profiles revealed some associations between species in different populations and probably vertical transmission of *E. coli* and *K. pneumoniae* between pregnant women and neonates or horizontal transmission between pregnant women, neonates, and healthcare workers.

The ERIC profiles of *E. coli* isolates revealed that in cluster E4 a pair of mother-neonate isolates (PO264B - N2264B) collected on the same date having 100% similarity although they displayed discordant MDR phenotype. Furthermore, this mother-neonate isolate pair shared a common ancestor with isolates from another mother (PO231A) and two other neonates (N2220 and N2234) collected seven days before this pair (Figure 4). In contrast, in cluster E7, a 100% similarity was observed between isolates originating from a mother (PO279C) and a neonate born to another mother (N2292) with a four-day collection lapse and both being MDR. These discordant mother-neonate isolates further shared a common ancestor with a caregiver (HW002A) collected 48 hours before the neonate and 48 hours after the mother (Figure 4). Intriguingly, a triplet of neonatal isolates (N2249A, N2247, N2246) shared 100% similarity with a maternal isolate (PO284A) collected one week apart and had a common ancestor with a neonatal isolate (N2235) collected two days before the neonatal isolates. Besides, the mother-neonate pair relatedness, *E. coli* isolates from mother-mother and neonate-neonate pairs sharing 100% similarity were observed in cluster E9-E13 (Figure 4).

**Figure 4:**
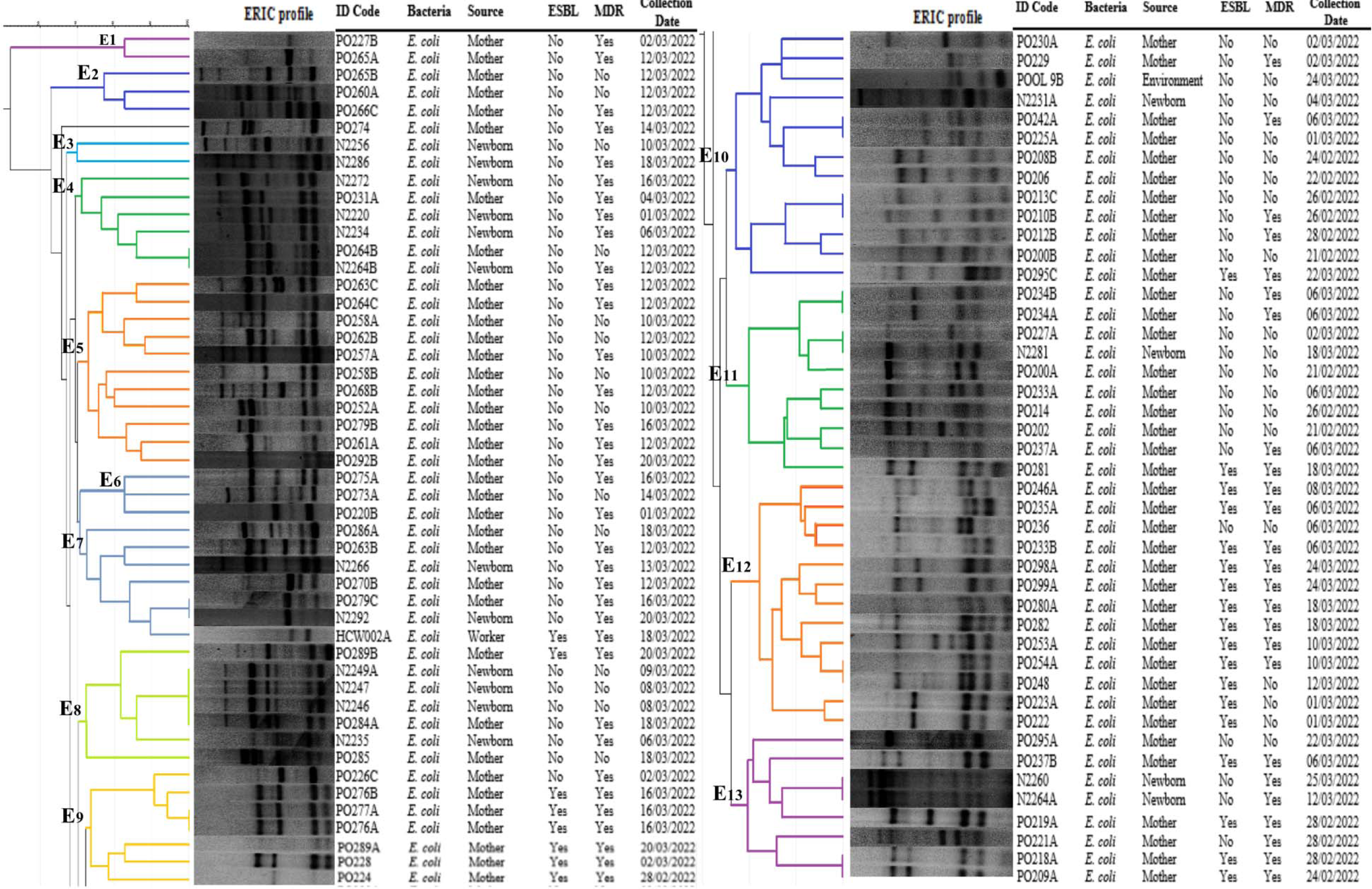
Genotypic relationship of *E. coli* strains (n = 93) detected from mothers, neonates, healthcare workers and the environment. Dendogram generated by GelJ using UPGMA method and the Dice similarity coefficient.

The ERIC profiles of *K. pneumoniae* revealed that none of the neonatal *K. pneumoniae* isolates were linked to the respective maternal isolates. Such discordance was observed for the neonatal isolates NN2223B (K6), NN2290 (K7), N2278 (K9) and N22218B (K10) sharing common ancestors with other isolates from other mothers, PO266A (K6), PO290 (K7), PO284C (K9) and PO299B. Interestingly, NN2276 (K9) shared a common ancestor with an isolate detected from a worker (HCW008A) (Figure 5).

**Figure 5:**
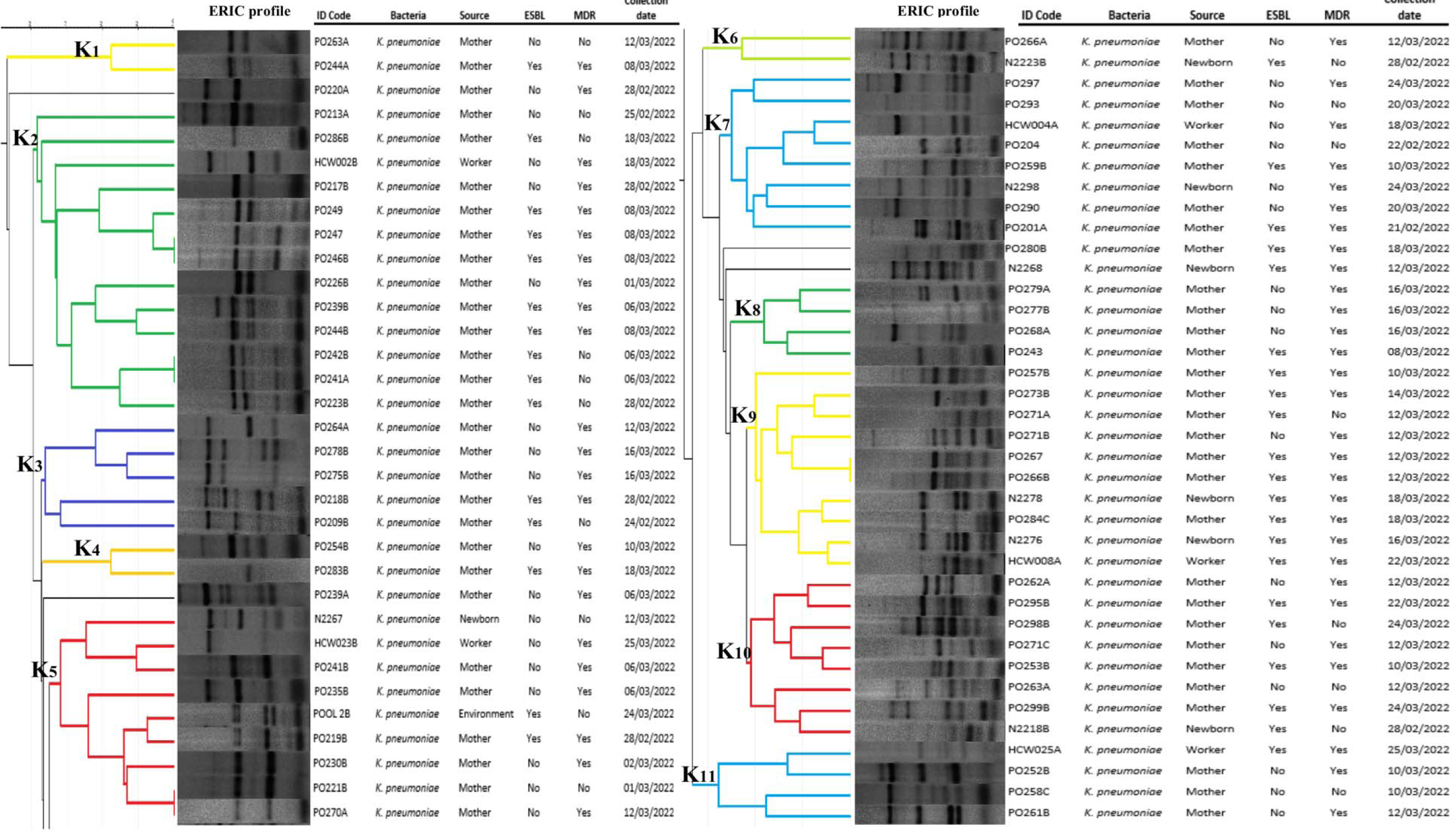
Genotypic relationship of K. pneumoniae isolates (n = 71) detected from mothers, neonates, healthcare workers and the environment. Dendogram generated by GelJ using UPGMA method and the Dice similarity coefficient.

## DISCUSSION

There is limited information about the burden and transmission of resistant bacteria in mothers and neonates in Cameroon, yet these are critical to implementing tailored prevention measures and achieving the United Nations Sustainable Development Goal (SDG) target 3.2.2. that aims for countries to have ≤12 neonatal deaths/1000 live births by 2030 (3, 4). Cameroon needs to increase its efforts by three to five times to expect to achieve this target by 2030 (1).

This study that aimed to determine the phenotypic and molecular features of *E. coli* and *K. pneumoniae* in a labour ward in Yaoundé, Cameroon revealed an elevated prevalence (rates ranging from 20-87%) of colonization by *E. coli* and/or *K. pneumoniae*. One of the most striking findings of this study was the high prevalence of MDR in pregnant women (90%, 73/81) and neonates (91%, 19/22). This unexpected, elevated prevalence of MDR can be explained by the absence of antimicrobial stewardship policies that favours irrational use of antibiotics in the human health sector, self-medication and over the counter supply of antibiotics. Additionally, the suboptimal water, sanitation, and hygiene (WASH), poor or non-existence of infection prevention and control (IPC) measures programme in the country and healthcare settings, coupled with absence of surveillance and monitoring system contribute to facilitate the emergence and dissemination of resistant bacteria. The detection of MDR-*E. coli* and MDR-*K. pneumoniae* might worsen the prognosis of neonates in case of infections.

MDR-*E. coli* and MDR-*K. pneumoniae* have often been involved in life-threatening neonatal infections, particularly sepsis, and were identified as the leading neonatal killers in 2019 (5). There is limited available data from Africa, our findings are similar to those of a Ghanaian study where 99% of Gram-negative bacilli growth was recorded in hospital environments, with over 80% of ESBL-producing isolates originating from the obstetrics and gynecology wards (15). However, the respective prevalences of *E. coli* (55%) and *K. pneumoniae* (45%) in pregnant women in our study are higher than those obtained in a study investigating vaginosis in pregnant women in Ethiopia with 25% *E. coli* and 2.3% *K. pneumoniae* (16). These differences may be explained by the collection methods used, the type of sample, and the different geographical locations.

On the other hand, the respective prevalence of *E. coli* and/or *K. pneumoniae* in neonates were 73% and 27%. These results are contradictory to those obtained in a study assessing the prevalence and risk factors of antimicrobial resistance in neonatal sepsis in Ethiopia where *K. pneumoniae* (79%) was the leading Gram-negative bacteria followed by *E. coli* (8%) (16). This difference may be explained by the follow-up period of neonates until 60 days after birth in the Ethiopian study. Interestingly, 91% (20/22) of the positive neonates were born to positive mothers, 27% and 18% of whom born to *E. coli* and *K. pneumoniae* positive mothers suggesting a probable vertical transmission from mothers to their neonates. This finding is in staggering contrast with a well-powered systematic review conducted in Africa that revealed a 27% prevalence of MDR-Gram-negative bacteria transmission from mothers to neonates, and 19% prevalence of neonatal ESBL-*Enterobacterales* colonization (9). The scarcity of studies from Central African countries in this systematic review could explain this discrepancy.

Of great concern, was the high prevalence of MDR-ESBL *K. pneumoniae* among neonates (100%), healthcare workers (83%) and mothers (60%). The fact that most healthcare workers admitted washing their hands between five to ten times a day and only after a medical act on a mother or a neonate gives credence to the hypothesis that sub-optimal IPC could contribute to the persistence and dissemination of MDR-ESBL-*K. pneumoniae* in the environment and subsequently to mothers and neonates. Acknowledging that hand hygiene has a significant role in reducing bacterial infections in healthcare settings when implemented adequately and timely, the World Health Organisation developed the five moments for hand hygiene guideline (17). In this guideline, the first two moments for hand hygiene that are essential to prevent the transmission of germs in hospital are before touching a patient (moment 1) and before performing a procedure on a patient (moment 2) (17). *K. pneumoniae* which is capable of resisting on healthcare workers’ hands or in the environment for a long period of time, could easily spread from healthcare workers or the environment to mothers or neonates when hand hygiene or hospital disinfection is not well implemented especially for the remaining moments, after a procedure (moment 3), after touching a patient (moment 4) and after touching patient surrounding (moment 5) (17, 18).

This can further explain the fact that one worker despite admitting washing his/her hand more than 30 times per day was colonised by MDR-ESBL-producing *K. pneumoniae*. It is thus plausible to surmise that healthcare workers by neglecting the moments 3-5 contribute inadvertently to the contamination of hospital environment and subsequent spread of resistant bacteria to mothers and neonates. This is particularly true for *K. pneumoniae* compared to *E. coli* given the presence of capsular membrane that ensure its survival in extreme environmental conditions. Smit et al. (2018) demonstrated that ESBL-*K. pneumoniae* were transmitted between humans and 22% of neonate infections were due to environmental source (19). Furthermore, Dramowski et al. (2022) emphasized the importance of WASH and IPC in the early and heavy bacterial colonization of neonates in resource-limited settings and recommended sustained neonatal IPC and surveillance programs in neonatal units in these settings (20). Healthcare workers colonized with MDR-ESBL isolates could thus contribute to the ongoing transmission of resistant bacteria to neonates where they could subsequently cause life-threatening neonatal infections such as sepsis, meningitidis or pneumoniae.

Infections caused by ESBL-producing *E. coli* and *K. pneumoniae* may be co-resistant to many other classes of antibiotics, resulting in the emergence of so-called multidrug-resistant bacteria (resistance to at least one antibiotic from three or more antibiotic families) (21). Both ESBL production and MDR is threatening considerably the management of neonatal infections and the prognosis of neonates (22). In this study, *bla_CTX-M_*, *bla_TEM_*, and *bla_SHV_* were widely detected at an overall prevalence of 75%, 43% and 42%, respectively. The *bla_CTX-M_*gene was most frequently isolated in women (81.35%) while *bla_CTX_*_-*M*_ and *bla_TEM_* co-dominated among neonates and healthcare workers. These results were different from those found in Sudan where the predominant gene was *bla*_TEM_ (86%) (23). This difference could be due to the abusive and inappropriate use of β-lactam antibiotics, especially cefotaxime in Cameroon.

ERIC genotypes revealed significant associations between isolates from pregnant women, neonates, and healthcare workers for both *E. coli* and *K. pneumoniae*. Only one dyad of mother-neonate *E. coli* isolates (PO264B–N2264B) was detected with 100% similarity, and these shared a common ancestor with a batch of isolates originating from a mother (PO231A) and two other neonates (N2220 and N2234) collected seven days apart. Most alarming was that all these neonatal isolates of this cluster were MDR-non-ESBL producers while maternal isolates were neither MDR nor ESBL. These results call into question the hand hygiene of the nursing staff and the environment in the delivery room. In fact, the discrepancies observed between neonatal and maternal isolates may be due to either temporary or permanent hand colonization of worker by *E. coli* or *K. pneumoniae* upon a contact with colonized neonates or mothers and the contaminated hospital environment.

The 100% similarity between two MDR-ESBL-*E. coli* isolates from two mothers collected on the same date raises the issue about the quality of the maternity ward environment, especially that of the labour room. It reveals the fluid transmission of bacteria within this maternity ward and highlights the limited prevention control measures notably hand hygiene existing in this hospital ward where bacteria can spread from one neonate to another or from one mother to another neonate horizontally via healthcare workers. Although we did not find significant bacterial loads in the environment in this study, it is important to note that the environment cannot be overlooked as a risk factor for contamination of mothers, neonates, and healthcare workers.

Notwithstanding, this study has several limitations. First, the limited sample size precludes any robust conclusion on the real burden of MDR and ESBL-*E. coli* and *K. pneumoniae* in all sources at the country level. Second, neonates could not be followed up to assess the outcome of this carriage by these resistant bacteria. Third, we could not identify the source of contamination nor ascertain the mode (vertical vs hospital-acquired), route of transmission especially in mothers and neonates. Fourth, it was not possible to investigate all resistance genes encoding for ESBL and MDR production. Finally, by investigating only the labour room, it is impossible to know if the carriage status of mothers was the same throughout the pregnancy. Despite these limitations, the study contributes to fill significant data gaps on the burden and genetic characteristics of *E. coli* and *K. pneumoniae* in a vulnerable population in Cameroon.

## CONCLUSION

This study suggests that MDR- and ESBL-*E. coli* and *K. pneumoniae* are actively circulating in labour ward with high prevalence in this confessional hospital of Yaoundé. It urges the imperative for stringent infection prevention and control measures particularly hand hygiene, in conjunction with antimicrobial stewardship programmes in the country. It advocates for the implementation of routine screening for multidrug-resistant and/or ESBL-producing *Enterobacterales* at least among at-risk pregnant women and neonates, as these could contribute to save lives in case of life-threatening neonatal infections. More advanced genomic studies are required and should be implemented to fully elucidate the transmission routes and sources of these resistant bacteria in Cameroon.

## Data availability

All data generated during this study are included in the article.

## Acknowledgements

We are grateful to the participants who provided samples for analysis in this study.

## Author Contributions

**Conceptualization**: Axelle Njeuna, Luria Leslie Founou, Ariel Blocker, Stephen Bentley and Lucien Honoré Etame Sone.

**Data curation**: Axelle Njeuna, Patrice Landry Koudoum and Aurelia Djemako Mbossi

**Formal analysis**: Axelle Njeuna, Luria Leslie Founou

**Investigation**: Axelle Njeuna, Patrice Landry Koudoum and Aurelia Mbossi.

**Project administration**: Luria Leslie Founou, Raspail Carrel Founou, Lucien Etame Sone.

**Software**: Axelle Njeuna, Patrice Landry Koudoum and Luria Leslie Founou.

**Supervision**: Luria Leslie Founou, Raspail Carrel Founou, and Lucien Honoré Etame Sone.

**Writing – original draft**: Axelle Njeuna, Luria Leslie Founou

**Writing – review & editing**: Luria Leslie Founou, Raspail Carrel Founou, Stephen Bentley, Ariel Blocker, Lucien Honoré Etame

**Approval and validation**: All authors

## Supporting Information

**Table S1.** List of selected environmental sampling sites

