## Supplementary Table 1 for "Maternal and Neonatal Colonization with Multidrug Resistant and Extended Spectrum ß-Lactamase Producing *Escherichia coli* and *Klebsiella pneumoniae* in a Cameroonian Labour Ward"

**Supplementary Table S1**: List of selected environmental sampling sites

| Level of risk | Sampling sites | Different pools |
| --- | --- | --- |
| High contamination risk | Five randomly sampled cradles in different rooms | POOL 2 : Neonates suite beds |
|  | Take five random beds in different rooms | POOL 9 : Mother’s postpartum beds |
| Medium contamination risk | - Delivery bed 1 - Bed for labor and delivery 2SA - Bed in ICU 3 - Plastic curtain, - Grab bar | POOL 4 : Equipment in contact with pregnant woman in delivery room |
|  | - Bench top - Clamp - Delivery equipment table - Tensiometer - Thermometer | POOL 5 : Materials used on the pregnant woman |
|  | - Radiant - Baby Bed - Vitamin K Cabinet - The balance - The meter | POOL 6 : Baby equipment in the delivery room |
|  | - Labor Bed 1 - Labor Bed 2 - Labor Bed 3 - Wall - Gallows | POOL 3 : Labor Room |
| Low contamination risk | - Registration table - Register - Patient chair - Wall and window - Personal Bench | POOL 1 : Reception |
|  | - Exam Room Door - Delivery room door - Entrance door - Restroom door - Entrance Wall | POOL 8 Labor and Delivery Room Doors |
|  | Four random room entrance doors + the post-partum entrance door. | POOL 7 : post-partum room Doors |
|  | - Lockers - walls - windows - Bag tables - Toilet doors | POOL 10 : Locker Rooms |

High risk: High probability of contact and exchange between healthcare workers, pregnant women and neonates.

Medium risk: Medium contact or exchange between healthcare workers and/or pregnant women and/or neonates.

Low risk: Low contact or exchange between the three populations, and contact occurring with healthcare workers or pregnant women or neonates.
